# Air-Liquid Interface Induced Epithelial Delamination

**DOI:** 10.1101/2023.08.14.553291

**Authors:** Chunzi Liu, Gerald G. Fuller

**Affiliations:** Department of Chemical Engineering, Stanford University Stanford, CA 94305

## Abstract

Many epithelial tissues reside at air-liquid interfaces, as exemplified by the ocular epithelium, oral mucosa, and alveolar epithelium. The interfacial tension across the epithelial tissues imposes a mechanical challenge to the homeostasis of the tissue. However, the interplay between interfacial properties and homeostasis in biological samples has been overlooked due to a lack of suitable measurement methods and theoretical developments. Here we described a surprising observation in which the surface energy at cell-air interface is sufficient to delaminate a stratified ocular epithelium from its substrate. We demonstrated that the interfacial tension at the epithelium-fluid interfaces can be measured using a modified Schultz method. The measured value is conceptually and numerically distinctive to the tensile modulus measured by deformation-based methods, such as micropippeting and tissue surface tensiometers. Furthermore, a mechanical analysis at the cell-air-liquid triple line during the delamination process revealed a strain hardening behavior of the epithelial layers. Finally, perturbations on different junctional protein complexes revealed that a delicate balance among cortical tension, focal adhesion, and cell-liquid interfacial tension is required for the epithelial tissue mechanical stability.

## 1 Introduction

Epithelial tissues are compact cell layers that protect organs from environmental interrogation. Residing at organ interfaces, epithelial tissues encounter polarized mechanical environments between apical and basal sides, including varied viscoelasticities and fluid shear rates [1, 2, 3]. Owing to drastic environmental changes across epithelial tissues, the interfacial properties of epithelial cells are crucial in facilitating their barrier function and maintaining the tissue mechanical integrity. However, the interfacial properties of epithelial tissues and their interactions with organ interfaces have been largely overlooked.

Among organ interfaces, the air-liquid interface is unique in its potential of mechanical interaction with epithelial tissues due to its high interfacial tension. Many epithelial tissues are exposed to air-liquid interfaces, as exemplified by ocular epithelium, oral mucosa, and alveolar epithelium. A back-of-the-envelope estimation shows that the force exerted on a single cell by the surface tension at air-liquid interface is on the order of 1 *μ*N, comparable to the forces exerted by other endogenous molecular machineries such as E-cadherin junctions and focal adhesion complexes [4, 5], suggesting that surface tension potentially contributes to the mechanical stability and homeostasis of epithelial tissues. A loss in mechanical stability can cause severe tissue failure and subject the inner organs to environmental pathogens [6, 7]. However, tissue mechanical stability analysis has largely overlooked the contribution from epithelial tissue surface tension.

Interfacial tension reflects the nature and strength of intermolecular forces at the interfaces. When one phase at the interface is air, the interfacial tension is also known as the surface tension. For biological systems, surface tension is usually obtained from the Young-Laplace relation by correlating the applied pressure and the subsequent cell or tissue deformation, as exemplified by tissue surface tensiometery and micropipette aspiration [8, 9]. However, as cells or tissues are actively deformed during these measurements, the cell response is contributed mainly by the contractile forces provided by cytoskeletal networks. Therefore, tissue surface tension obtained via deformation-based methods does not reveal the interaction between the cell surface and the external fluid phase.

In addition to deformation-based methods, the contact angle at the cell-air interface is reported as a measure for molecular interaction strength at this interface, and therefore, indicates the interfacial energy at this interface. Previous contact angle measurement at the cell-cell interface provides the ratio between cell surface tension and cell-cell adhesive force [10, 11]. Herein we adapted the Schultz’s method [12, 13, 14], a conventional method to measure the high energy solid surface energy based on contact angle measurements, to measure the surface tension of epithelial layers and the interfacial tension at the cell-aqueous solution interface.

The ocular epithelium is naturally exposed to an air-liquid interface, and its surface tension is crucial in determining the constant hydration mechanism at the ocular surface. The human ocular epithelium is a stratified 6-9 layers of epithelium covering the corneal and scleral surfaces. Its interfacial properties are paramount in maintaining the tear film homeostasis and, subsequently, a sustainable lubrication of the ocular surface [15, 16, 17, 18, 19]. Measuring the ocular epithelium’s surface tension can illuminate the mechanism behind the tear film stability and the sustainable ocular surface hydration [20, 21]. However, measuring the exact surface tension of ocular epithelium is challenging due to its hydrophilic nature and complex molecular compositions. Herein we reported the surface energy of stratified corneal and conjunctival epithelia with a modified Schultz’s method.

The mechanical stress exerted by the air-liquid interface potentially alter the biological functions and the physiology of tissues *in vivo*. Previous literature has shown that imposing a surface tension can induce physiological changes in cerebral organoids and breast cancer cells via mechanotransduction pathways similarly regulated upon changes in substrate stiffness [22, 23]. Herein we describe a surprising phenomenon wherein the surface tension at the air-liquid interface is sufficient to delaminate ocular epithelial tissues from solid substrates. The delamination process enables a rheo-mechanical study of the ocular epithelium under extreme mechanical perturbations. Via regulations on epithelial cortical tension and cell-substrate adhesion, we reveal a phase diagram that governs the epithelial tissue mechanical stability at the air-liquid interface, including a substantial contribution from surface tension. The results illustrate that the mechanical stress near the air-liquid interface actively participates in tissue functions, potentially in mechanotransduction and cell decision making processes.

## 2 Results

### Air-liquid interface induces epithelial delamination

To study the mechanical interactions between the air-liquid interface and the ocular epithelium, we utilized a contact angle goniometer with a tilting stage in the captive bubble geometry as previously described [19, 17], in which the cell layers are immersed in an aqueous environment facing down and in contact with an air bubble. Stratified telomerase-immortalized human corneal epithelial (hTCEpi) and immortalized human conjunctival epithelial (HCjE) layers were chosen to represent the corneal and conjunctival epithelia as previously described [19, 17, 16]. The air-liquid interface was created by freely attaching an air bubble onto epithelial cell layers in an aqueous solution, as shown schematically in Fig 1A. With increasing tilting stage angles, *ϕ*, the normal stress and shear stress increase as *ρV g* cos *ϕ* and *ρV g* sin *ϕ*, respectively, where *ρ* is the density of the external fluid phase and *V* is the volume of the air bubble. Under the shear stress imposed by the air bubble, the epithelium exhibited three different dynamics, which we termed the static, delamination, and rupture behaviors.

**Figure 1.**
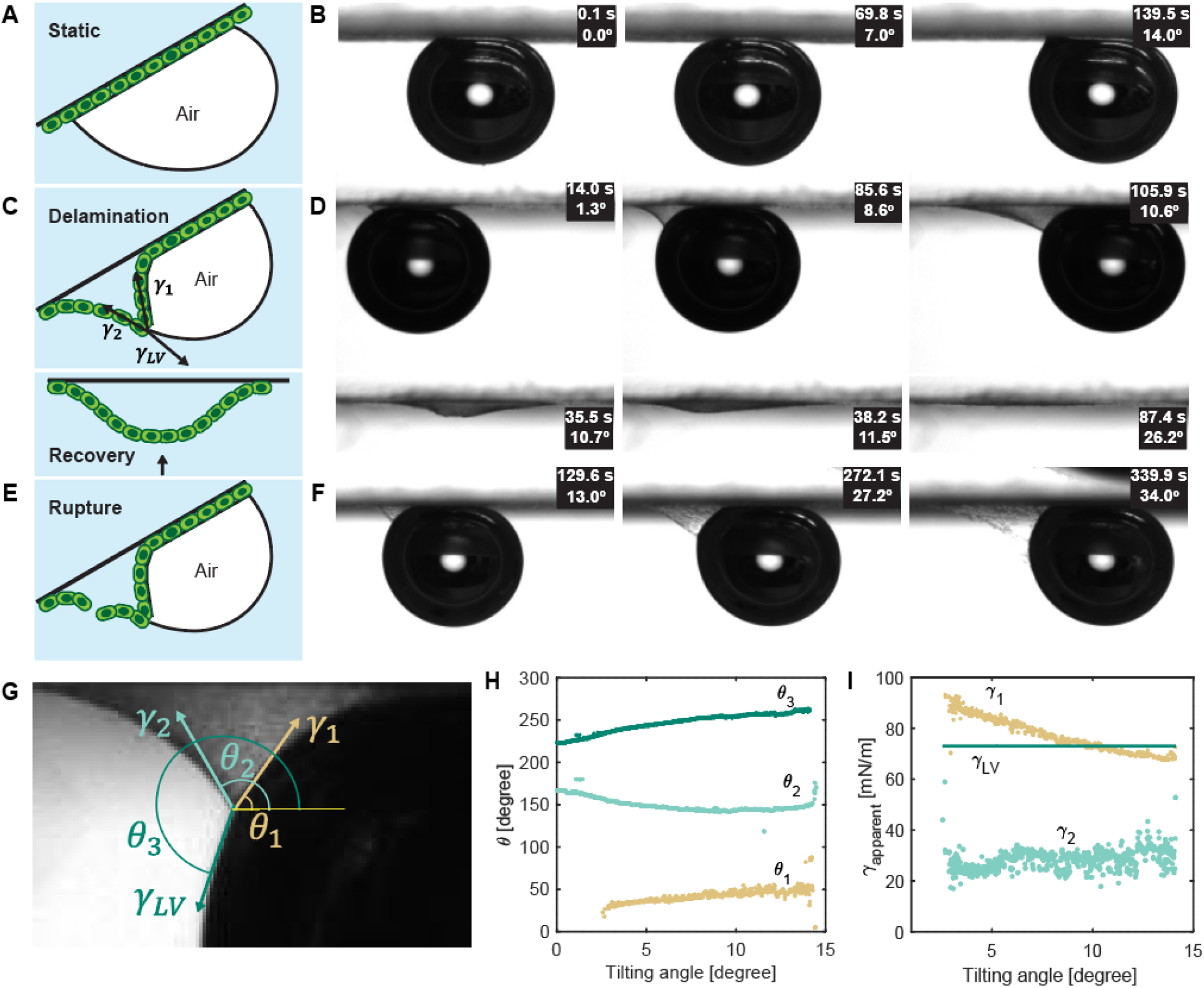
: Stratified ocular epithelium deformation in response to the air-liquid interface. Schematics and representative snapshots show three different behaviors: (A-B) static response, (C-D) delamination and recovery, and (E-F) rupture observed for stratified hTCEpi cell layers. G, schematic of stress analysis at the triple line during the delamination process. Quantification of relative angles (H) and apparent interfacial tensions (I) for three interfaces at the triple line. Surface tension of the aqueous phase, *γ*_*LV*_, was measured by pendant drop method for each experiment.

As shown in Fig 1B and Movie S1, the air bubble escaped from stratified corneal epithelial layers in phosphate buffer solution without inducing epithelium deformation, which is commonly seen in contact angle hysteresis measurements for solid substrates [19, 17]. We will refer to this behavior hereafter as the static behavior.

Surprisingly, corneal and conjunctival epithelial layers delaminated from the collagen-coated coverslip substrates after the cell-cell and cell-substrate adhesion was weakened with EDTA treatment as shown in Fig 1D and Movie S2. After the air bubble escaped, the delaminated epithelial layer recovered under the tensile stresses within the cell layer and reattached to the substrate. The aqueous phase in the experimental chamber transported through the epithelial layer via weakened cell-cell junctions. Visually, the delamination process resembles a three-fluid interface deformation [24]. However, a distinction should be drawn between these two phenomena. The deformation at fluid-fluid interfaces is driven by the viscous nature of the interface, while the epithelial layer deformation at air-liquid interface is largely driven by the elastic response contributed by the cytoskeletal networks, similar to the deformation induced at a fluid-fluid-solid interface as described previously [25, 26, 27, 28, 29].

In addition to static and delamination behaviors, rupture was also observed in stratified conjunctival layers immersed in phosphate buffer solution, as shown in Fig 1F and Movie S3. Initially, the epithelium delaminates from the substrate as observed in the case of EDTA-treated epithelium. As the stage tilting angle increases, the shear stress imposed by the air bubble overcomes the cohesive strength in the axial direction of the epithelial layer, which induces epithelial cell dissociation and layer rupture.

To further understand the stress conditions and the epithelial layer mechanical properties that distinguish these behaviors, we quantified the magnitudes of interfacial stresses as functions of the tilting angle, *ϕ*. Three nominal stresses contribute to the delamination process, the stress at the air-cell-liquid interface, *γ*_1_, the stress at the liquid-cell-liquid interface, *γ*_2_, and the surface tension at the air-liquid interface, *γ*_*LV*_, as schematically shown in Fig 1G. Recognizing that the time scale of the tilting motion (1 s) falls between the characteristic time scale of cell elastic response provided by the cytoskeletal contractile forces (1 ms) and epithelium rearrangement driven by cell motility (1 hr), we applied a force balance at the triple line, ***γ***_**1**_ + ***γ***_**2**_ + ***γ***_***LV***_ = **0**, which resembles the Neumann’s construction applied to the deformation at a three-fluid interface [30, 24]. To solve the force balance, *γ*_*LV*_ was determined by a pendant drop method (SI appendix, Methods and Fig S1) and the geometry of the triple line was quantified with a customized MATLAB code as shown in Fig 1H. The surface tension of the aqueous phase, *γ*_*LV*_, was measured to be (61.8 *±* 1.0) mN*/*m in the presence of corneal epithelial cells, and (64.1 *±* 0.5) mN*/*m in the presence of conjunctival epithelial cells. These values are lower than the surface tension of pristine PBS, 69.5 mN*/*m, likely due to the adsorption of secreted proteins from epithelial cells onto the air-liquid interface. By solving the force balance, the magnitudes of *γ*_1_ and *γ*_2_ as a function of the chamber tilting angle are reported in Fig 1I. A relaxation behavior was observed for the stress on the air-cell-liquid branch, *γ*_1_, while the stress on the liquid-cell-liquid branch, *γ*_2_, remained stable over the delamination process.

To understand the molecular origin of the nominal stresses at the liquid-cell-liquid and air-cell-liquid interfaces, we recognized that *γ*_1_ and *γ*_2_ have significant contributions from cell-liquid and cell-air interfacial tensions, while the residual stress arises from a strain-dependent response function, *γ*^*elastic*^(*ϵ*), where *ϵ* is a strain value that describes the epithelium deformation. This response function is an intrinsic property of the epithelial layer, which microscopically describes the cohesive strength generated by the cytoskeletal cortical tension and the cell-cell adhesive force. In comparison, the interfacial tensions at the cell-liquid interface, *γ*_*CL*_, and at cell-air interface, *γ*_*CV*_, arise from the intermolecular interactions between cell surface molecules and the external fluid phase (air/liquid) in the direction normal to the epithelial layer. Concisely, *γ*_1_ and *γ*_2_ are given by,

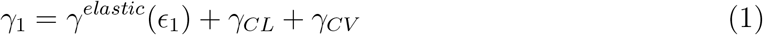

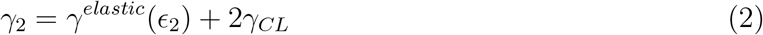

### Surface energy of stratified ocular epithelia

To obtain the response function of epithelial layers under extensional stresses, *γ*^*elastic*^(*ϵ*), the cell surface energy, *γ*_*CV*_, and the cell-liquid interfacial tension, *γ*_*CL*_ are required to be independently measured. However, measuring the surface energy of layers of living cells is technically challenging. Epithelial layers exhibit high surface energies due to the hydrophilic, negatively charged macromolecules present on cell surfaces, such as glycoproteins and polysaccharides [31, 32]. Therefore, surface energy measurements suitable for low energy solids, such as Zisman’s critical surface tension method and Neumman’s single contact angle measurement [33, 34, 24], are insufficient to determine the surface energy of epithelial layers. To measure the high surface energy of live epithelial layers, we utilized Schultz’s method with reversed liquid phases [12, 13], in which the contact angles of various non-polar liquid droplets on epithelial layers are measured in an aqueous phase as shown Fig 2A. The contact angles of live epithelial layers against different non-polar liquids with known surface energies are then converted to a surface energy value of the live epithelial layer via a linear regression process outlined in the Method section. Briefly, Fowkes’ expression for solid-fluid interfacial tension [14] and Young’s equation at solid-fluid-fluid triple line are combined to demonstrate a linear relation between 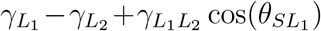 and 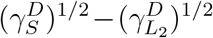, where the subscripts *L*_1_ refers to the non-polar liquid phase, *L*_2_ refers to the polar (aqueous) liquid phase, *S* refers tp the solid phase (epithelial layer), and the superscript *D* refers to the dispersive contribution of the interfacial tension. The slope from the linear regression, 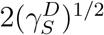, signifies the dispersive surface energy of the epithelial layer, and the intercept,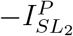, signifies the polar contribution to the interfacial energy at cell-aqueous interface.

**Figure 2.**
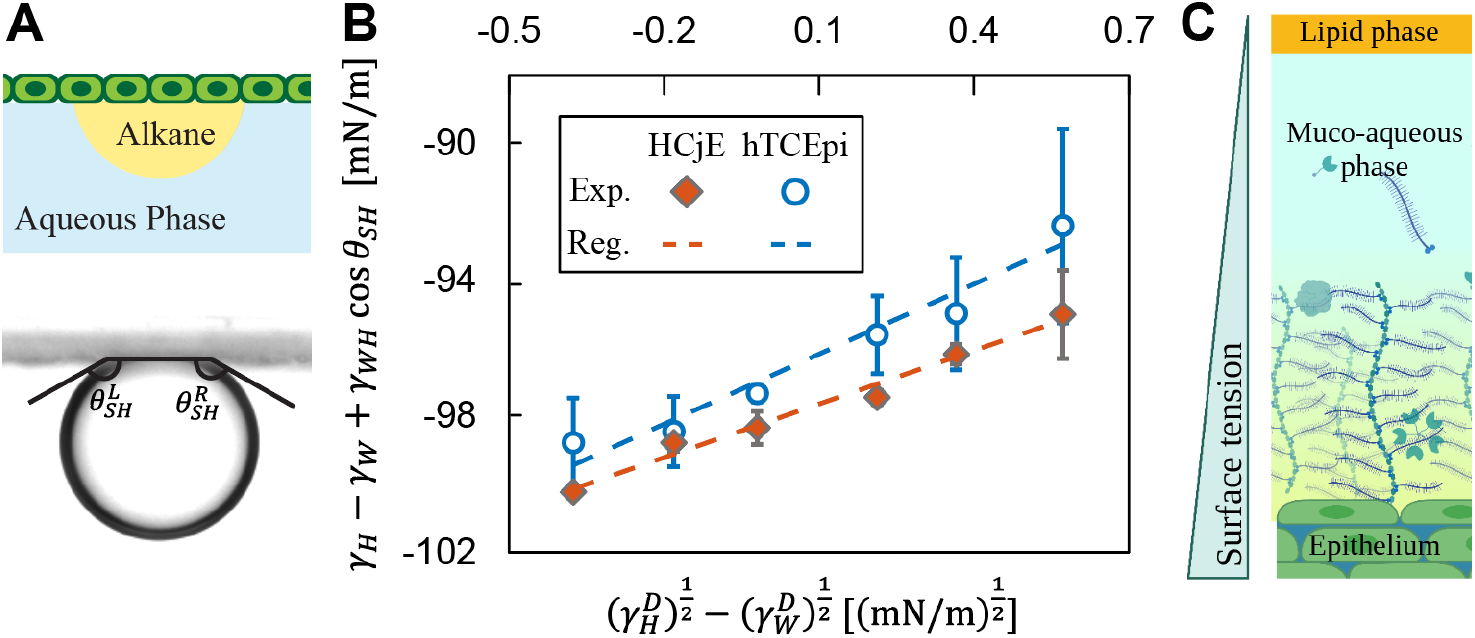
: Surface energy measurements of epithelial cells. (A) Schematic and representative image of contact angle measurements for a non-polar liquid droplet in an aqueous bulk phase on the cell surface. The average of left, 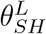, and right, 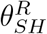, contact angles is used as the contact angle for a single measurement. (B) Linear regressions using contact angle data for stratified HCjE and hTCEpi cell layers. The error bars indicate standard deviations of a sample number *n* = 3 *−* 5 for each alkane/cell interface. A minimum of five contact angles were measured for each sample. (C) A schematic showing the surface tension gradient on the ocular surface.

For this study, six linear alkanes were chosen as the non-polar fluid phase. To measure the surface energy of corneal and conjunctival epithelial layers, the stratified epithelial layers on coverslips were immersed in an aqueous phase and an alkane droplet was generated to be attached onto the epithelial layer. The contact angles of the alkane droplets were quantified with a customized image analysis program and reported in Fig 2B. The parameters, including the surface tensions of alkane and aqueous phases, are reported in SI Appendix Table S1. From linear regression done on Fig 2B, the surface energies of stratified corneal and conjunctival epithelial cells, *γ*_*S*_, were measured as (56.0 ± 1.5) mN*/*m and (54.2 ± 1.0) mN*/*m, respectively, as reported in Table 1. From Table 1, the two cell lines exhibit similar surface energies that are predominantly contributed by the polar interactions at the cell-fluid interface (large 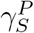). Both epithelial layers exhibit diminishing interfacial tensions against PBS (small *γ*_*SW*_), in agreement with the high mucin and glycoprotein expressions on their cell membranes [19, 17, 16]. As the cell layers are not actively deformed during alkane contact angle measurements, the surface energy reported here strictly reflects the intermolecular interactions at the cell-fluid interface. This distinguishes our cell surface energy values from the surface tension values acquired with tissue surface tensiometers and micropipette aspirations that are largely contributed by the epithelial layer cortical tension in response to imposed stress or deformation [8, 9].

**Table 1:**
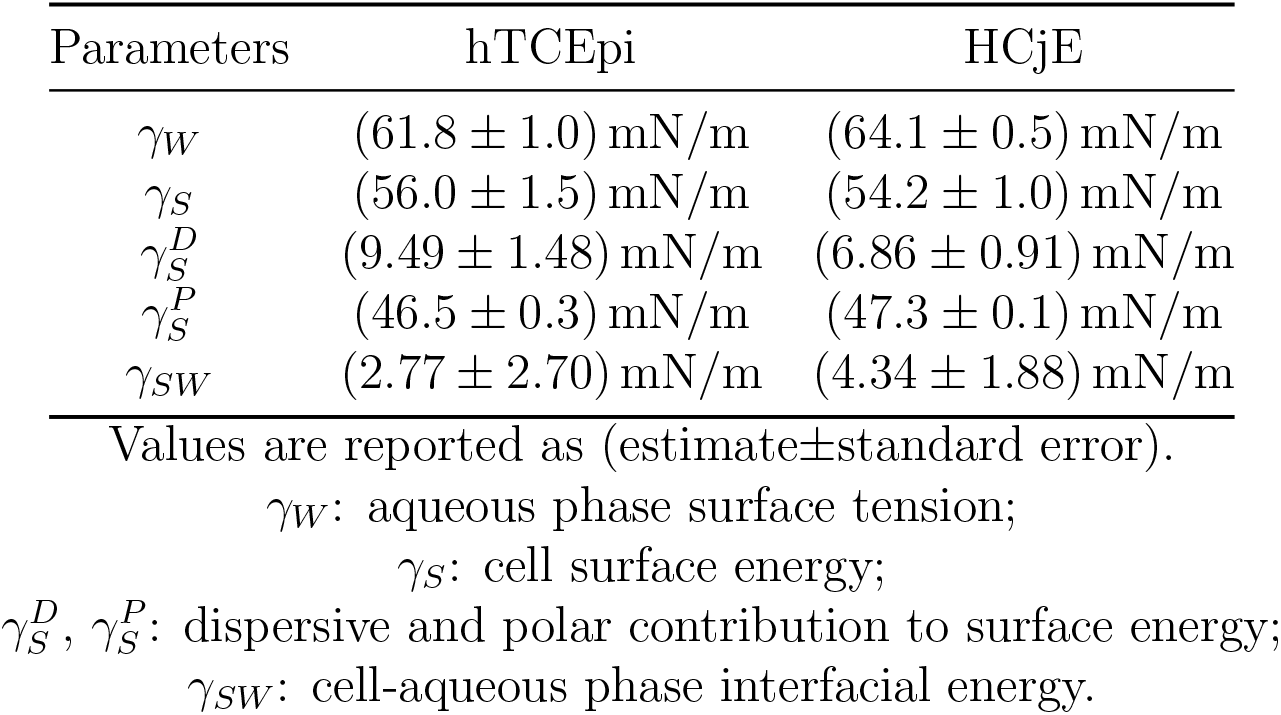
Surface energies of stratified hTCEpi and HCjE layers measured by modified Schultz’s method.

### Epithelium yielding behavior under large deformation

To study the rheological behaviors of ocular epithelium under extensional stresses, we correlate the in-plane stress exerted on the epithelium with its axial deformation. The inplane stress exerted on epithelia was obtained by subtracting the contributions of cell-air and cell-liquid interfacial tensions from the apparent stress, namely, *γ*^*elastic*^(*ϵ*) = *γ*_*i*_ *− γ*_*interface*_. An engineering strain was used to describe the deformation of the epithelial layer, in which the strain, *ϵ*, is defined as *ϵ* = (*L − L*_0_)*/L*_0_, where *L*_0_ and *L* are the epithelial layer lengths in projection before and after delamination, respectively. A schematic of the strain quantification is shown in Fig 3A. As the strain increases over the course of an experiment, *γ*^*elastic*^ also gradually increases as shown by a representation graph in Fig 3B. As the experimental chamber was tilting at a sufficiently low speed, we assume that the ocular epithelium deformation is in an equilibrium state. To obtain the response function of ocular epithelium subject to the air-liquid interfacial tension, *γ*^*elastic*^ is plotted as a function of epithelial deformation, *ϵ*, for hTCEpi epithelium under EDTA treatment in Fig 3C. The response function from different experiments collapsed onto a master curve, indicating that the slow-tilting assumption is valid.

**Figure 3.**
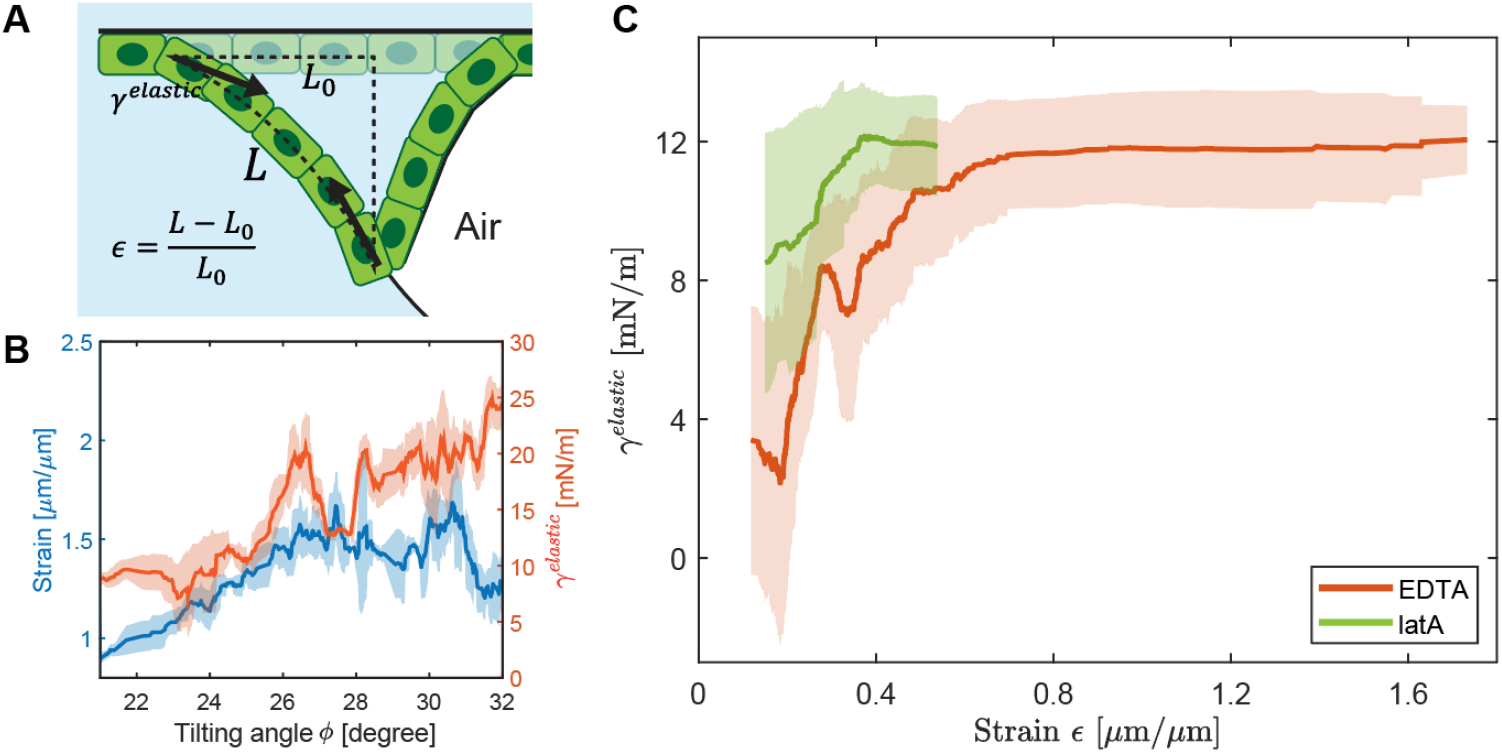
: Ocular epithelia showed yielding behavior under extreme deformations induced by the air-liquid interface. (A) A schematic showing the definition of the engineering strain, *ϵ*. (B) A representative graph showing *ϵ* and *γ*^*elastic*^ as a function of the tilting angle in a single delamination experiment. (C) A mastercurve between *ϵ* and *γ*^*elastic*^ for hTCEpi cell line under EDTA (*n* = 4) and latrunculin-A (*n* = 2) treatments. The central solid line indicates the moving average and the shaded region indicates the standard deviation.

From Fig 3C, the epithelium followed an elastic behavior at small strain values (<0.4), indicating that cytoskeletal contractile forces dominate the rheological behvaiors in this regime. The elastic behavior at small strains allowed us to estimate Young’s modulus for ocular epithelium. To estimate Young’s modulus, a linear regression was done on the stress-strain curve at small strain values. The slope is (27.2 ± 3.6) mN/m for hTCEpi cells under EDTA treatment. This value is higher than the previously reported epithelial monolayer tension (*O*(1 mN/m)) potentially due to differences in measurement geometry and epithelium geometry[35]. To compare the 2D modulus reported from the stress-strain curve here to the epithelial elasticity measured by atomic force microscopy, we estimate the epithelial Young’s modulus by dividing the slope by a cellular length scale of 10 *μ*m to obtain a modulus value around 2.7 kPa. This estimated modulus value is consistent with the Young’s modulus of 5 kPa measured for this cell line with atomic force microscopy [17]. To test the contribution of cortical tension to the epithelium elasticity, the cytoskeletal network is weakened with latrunculin treatment. After the latrunculin treatment, the response function slope at small strain is (24.3 ± 5.1) mN/m (Fig 3C), comparable to the value obtained under the EDTA treatment. Previous micropipette aspiration experiments indicate that cortical tension contributes significantly to epithelium elasticity. While exhibiting a similar slope in the delamination experiment, the response function for latrunculin-treated epithelial layers has a much lower maximum strain when the air bubble escapes compared to the EDTA-treated epithelial layers, corresponding to less apparent plasticity in the latrunculin-treated epithelial layers. Complementary to previous reports, the lower maximum strain at break for latrunculin-treated epithelial layers indicates that epithelial plasticity is largely contributed by the remodeling of cytoskeletal networks.

At large strain values (*>* 0.6), the response function exhibits a plateau region where the in-place stress remains constant while the epithelium continues to deform, phenomenologically similar to the plastic deformation experienced by metals and polymeric materials at large strains. However, under our experimental conditions, the EDTA-treated epithelia always exhibit full recovery after the air bubble is removed, differentiating the epithelial stress plateau behavior at large strains from conventional plastic deformation. Irreversible deformation was observed in latA-treated epithelia (Movie S4). As EDTA weakens the cell-substrate and cell-cell adhesion complexes while latA weakens the cytoskeletal network, these observations suggest that cytoskeletal network can rearrange under extreme deformation, giving rise to the reversible deformation in the plateau region. Once the ability to rearrange is impaired in the cytoskeletal network, the large deformation becomes irreversible.

### Epithelium mechanical stability

To understand the molecular origin of the delamination behaviors at the air-ocular epithelium interface, we prescribed a mechanical stability analysis on the system and developed an analytical phase diagram describing transitions among static, delamination, and rupture behaviors. Three stresses are hypothesized to maintain the mechanical stability of the ocular epithelium, which are the stress parallel to the epithelial layer based on cell connectivity, *γ*_*CC*_, the cell-substrate adhesion via focal adhesion complexes, *γ*_*CS*_, and the cell-aqueous phase interfacial tension, *γ*_*CL*_. The epithelium in-plane stress is contributed by the contractile cortical tension generated by the cytoskeletal network, while the cell-cell junctional adhesive strength dictates the maximum stress in parallel to the cell layer that does not induce epithelial cell dissociation. Under our experimental conditions, weakening cell-cell adhesion on corneal epithelium via EGTA treatment did not induce a change in epithelial behaviors, which indicates that cortical tension dominates the epithelial stability, in agreement with previous observations of cell mechanical study via micropipette aspiration and tissue surface tensiometery [11, 36].

To develop an analytical phase diagram for cell-air interactions, we prescribed stress conditions for adhesive and cohesive failures for the epithelial layers. As shown schematically in Fig 4A, an adhesive failure describes the process in which the normal stress on a single cell exceeds the maximum cell-substrate adhesion provided by focal adhesion complexes and cells are subsequently detached from the substrate, or *γ*_*LV*_ sin(*θ*) *> γ*_*CS*_, where *θ* is the contact angle at cell-air-liquid interface. As shown schematically in Fig 4B and observed in Movie S3, a cohesive failure describes the process in which the in-plane stress exceeds the maximum cell-cell junctional adhesion and the cell layer dissociates from the junctional sites, or 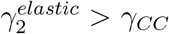, where *γ*^*elastic*^ is the in-plane elastic stress, and *γ*_*CC*_ is the maximum cell-cell junctional stress. In the following analysis, *γ*_*CC*_ is approximated by the maximum stress value on the stress-strain curve, or the ultimate tensile strength within the epithelial layer, *γ*^*UT S*^. We recognize that the in-plane elastic stresses in the attached and detached epithelial branches are the same, or 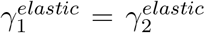, as the cell at the basal junction can only transmit the same force. Combining with geometric confinements, the cohesive failure condition is 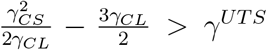. Two dimensionless stresses emerge from the analysis, which are the normalized maximum cortical tension, 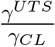, and the normalized maximum cell-substrate tension, 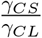. Rearranging the adhesive failure condition, the phase boundaries prescribed by adhesive and cohesive failures are,

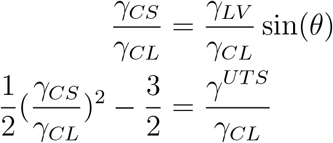

From the equations of state as shown in Eq **??**, we constructed phase diagrams for stratified corneal and conjunctival epithelial layers as shown in Fig 4C, in which *γ*_*LV*_ and *γ*_*CL*_ are reported in Table 1. The contact angle, *θ*_hTCEpi_, for stratified corneal epithelial layer was reported as 155.6° ± 1.1°in a previous literature [17], and *θ*_HCjE_ was measured to be 132.1° ± 1.3°. The delamination-rupture phase boundary (cohesive failure) is the same for both cell lines while the rupture-static phase boundary (adhesive failure) is determined by the relative strengths of liquid-vapor and liquid-cell interfacial tensions.

**Figure 4.**
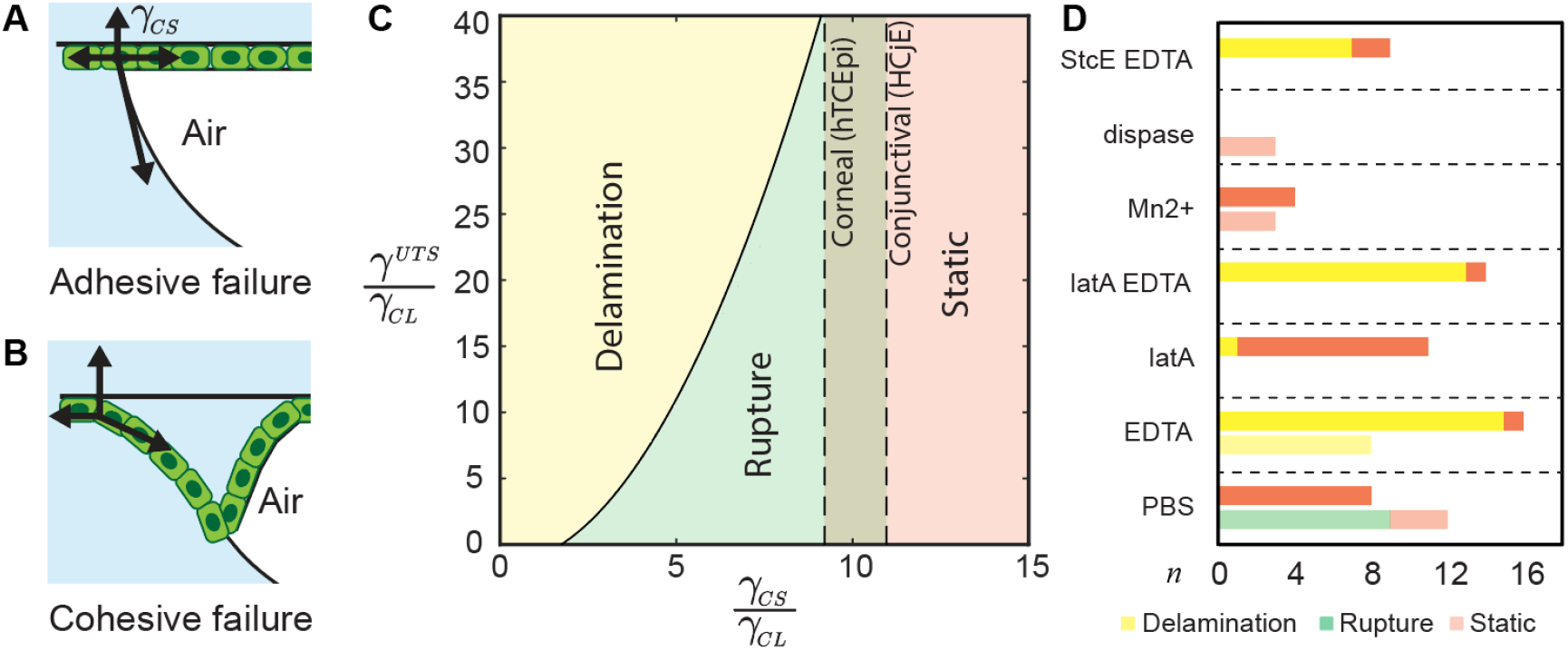
: Conditions to induce static, delamiantion, and rupture behaviors in epithelial tissues against an air-liquid interface. Schematics of A. adhesive and B. cohesive failure conditions. C. Phase diagram of epithelium mechanical behaviors against an air-liquid interface. The relative strength of cell ultimate tensile strength (a function of cell cortical tension and cell-cell adhesive force), cell-substrate adhesive force, and cell surface tension determines the epithelial mechanical behaviors. D. Pivot chart of ocular epithelium behaviors at air-liquid interface after various drug treatments. Data for hTCEpi is shown in the darker columns, HCjE is shown in the lighter columns. The experiment number, *n*, indicate the number of biological replicates. Behaviors were evaluated after 30 minutes in the aqueous phase.

To test whether the analytical phase diagram can predict epithelial behaviors in contact with the air-liquid interface, we targeted cortical tension and cell-substrate adhesive strength in stratified corneal and conjunctival epithelium using drug treatments. The epithelial layer behaviors after drug treatments are summarized in Fig 4D. Under control conditions, stratified corneal epithelium exhibited static behavior, and stratified conjunctival epithelium exhibited rupture behaviors. After the cell-substrate adhesion was decreased with EDTA treatment, both stratified corneal and conjunctival epithelial layers transitioned into delamination behaviors, agreeing with the adhesive failure at the low *γ*_*CS*_*/γ*_*CL*_ regime. The dispase treatment did not induce a similar transition potentially due to the spatially inhomogeneous transport of dispase onto the epithelial substrate (see Discussion). Increased cell-substrate adhesion via Mn^2+^ treatment transitioned conjunctival epithelial layers from rupture to static behavior, agreeing with the static behaviors at the high *γ*_*CS*_*/γ*_*CL*_ regime. Collectively, the phase transition via cell-substrate adhesion regulation agrees with the analytical phase diagram. Weakening cortical tension with latrunculin-A treatment did not induce a transition in stratified corneal epithelial layers, agreeing with agnostic nature of the static phase to the in-place stress. A double treatment of latrunculin-A and EDTA on stratified corneal epithelial layers did not induce a delamination-rupture phase transition, likely due to the extremely low *γ*_*CS*_ after the EDTA treatment. An attempt to modify the cell-aqueous phase interfacial tension was done with a mucinase treatment. However, no change in epithelial layer behavior was observed, as it has been shown that a lack of mucin contributes little to the epithelial tissue contact angle, which is a reflection of its surface energy [17]. More effective treatments are needed to alter *γ*_*CL*_ for epithelial layers.

## 3 Discussion

The high surface tension of ocular epithelium suggests that the tear film stability is maintained via a surface tension gradient along the ocular radial direction. Tear film is a thin layer of multi-component fluids that lubricates the spontaneous blinking motion and protects the ocular surface from pathogen adhesion. Its function depends on its complete wetting over the ocular surface, which is difficult due to a three orders of magnitude difference between the thickness of the tear film (*O*(10 *μ*m)) and the diameter of the ocular surface (*O*(1 cm)). Tear film is composed of a lipid layer and a muco-aqueous layer. Previous studies report that purified lipid phase and aqueous phase from the ocular surface have surface tensions of 46 mN/m and (53.2 ± 0.2) mN/m, respectively [37], higher than the ocular epithelium surface tension reported here. As schematically shown in Fig 2C, the surface tension gradient established by epithelium, aqueous, and lipid phases minimize the free energy along the radial direction of the ocular surface and, therefore, stabilize the tear film. A disrupted tear film exposes components of higher surface tension (aqueous phase and epithelium), driving the restoration of the established surface tension gradient. This simple argument does not account for the dynamical environment of the ocular surface due to the constant blinking motion. Further simulation efforts on the hydrodynamics of the ocular surface can help illustrate whether the surface tension gradient alone is sufficient to restore the tear film stability on an adequate timescale.

A homogeneous mass transport across the ocular epithelium is necessary for the delamination behavior in various drug treatments. While lowering the cell-substrate adhesion via EDTA treatment induces the delamination behavior, an enzymatic cleavage of extracellular components via dispase treatment did not induce a similar transition potentially due to a spatially inhomogeneous transport of dispase onto the epithelial substrate. A mature epithelium exhibits barrier function that does not permeate small molecules across epithelial cell tight junctions. Therefore, dispase is only delivered from the substrate boundaries that likely result in a whole cell sheet delamination prior to experiments. In comparison, EDTA weakens the cell-cell tight junction, and therefore, allows a homogenous transport and decrease of cell-substrate adhesion, in agreement with the observation that epithelia treated with dispase were observed to spontaneously delaminate from the coverslip edges. The mass transport across the epithelium is also important for the delamination dynamics as the aqeusou phase needs to transport via the epithelium with impaired cell-cell junctions, a process that can be modeled as fluid transport across a porous medium.

## 4 Conclusion

Here we have illustrated that the air-liquid interface can induce significant mechanical stresses in ocular epithelium, suggesting that the surface tensions should be taken into consideration for future biomechanical studies in tissues and organisms that encounter drastic environmental changes, including the processes of embryogenesis, organ transplant, pathogenesis, and microorganisms physiology. The epithelium surface tension measurement method introduced here is widely applicable to biological surfaces with a flat geometry. The measured ocular epithelium surface tension, combined with previous studies on tear film components, suggests that a surface tension gradient might contribute to the tear film stability. Further studies are needed for the delamination phenomenon in epithelial tissues that encounters spatially drastic changes, e.g., aveolar epithelium, gastrointestinal epithelium, and oral epithelium.

### Supporting Information Appendix (SI)

Supplementary methods, figures, tables, and movies are available in a separate SI Appendix file.

## 5 Methods

### Cell culture

Human telomerase reverse transcriptase-immortalized corneal epithelial cells (hTCEpi) and human immortalized conjunctival epithelial cells (HCjE) were obtained via previously described methods [19, 17, 16]. hTCEpi cells were used between passages 56 and 70. HCjE cells were used between passages 6 and 20. hTCEpi and HCjE cells were cultured in the growth medium (GM) composed of Epilife supplemented with Human Corneal Growth Supplements (HCGS) and 1% penicillin-streptomycin at 37 °C and 5% CO_2_. To induce differentiation and stratification, the GM was replaced by the stratification medium containing Dulbecco modified Eagle/F12 medium, 10% fetal bovine serum, 10 ng mL^*−*1^ EGF, and 1% penicillin-streptomycin.

### Delamination

The experimental procedure is similar to the contact angle hysteresis measurements described in a previous literature [17]. The method is briefly described here. The delamination experiments were conducted with a Ramé-Hart 290 contact angle goniometer. The captive bubble geometry was enabled with a customized acrylic chamber and a 3D-printed inset with indents for 18 mm square coverslips. Cells were grown on acid-washed coverslips and stratified for seven days prior to experiments. An inverted needle (Gauge No.25) was attached to the air dispenser. The experimental chamber was filled with PBS immediately after an oxygen plasma treatment. The coverslips with cell culture were gently washed with PBS for three times before being loaded onto the 3D-printed inset with the cell side facing down. The surface tension of the aqueous phase was measured before each experiment with the pendant drop method. During an experiment, an air bubble of 5 μL was created and freely attached against the cell surface. A video of the air bubble and the cell layer was recorded with the DROPimage Advanced software while the stage was tilting at 0.1 ° s^−1^.

### Surface energy of epithelial cell layer

A modified Schultz’s method was applied to measure the surface energy of epithelial cell surface, in which the contact angle of a non-polar liquid droplet on a cell layer is measured in an aqueous bulk phase. Hexane, heptane, octane, decane, dodecane, and heaxadecane were chosen as the non-polar liquid phase, and PBS was chosen as the aqueous bulk phase to avoid aging of the fluid interface driven by protein adsorption. An average of at least five contact angles were reported for each epithelial cell layer. The contact angles were quantified with a MATLAB code. Only one type of *n*-alkane liquid was chosen for an epithelial cell layer to prevent cross-contamination. At least three samples were measured for each cell/alkane pair.

To obtain surface energy from contact angle measurements, a brief derivation is described below for the geometry in Fig 2. For a non-polar, low density liquid droplet, *L*_1_, attached onto a cell surface, *S*, immersed in an aqueous phase *L*_2_, the interfacial tension at cell/non-polar liquid and cell/aqueous phase interfaces are given by Fowkes’ expression,

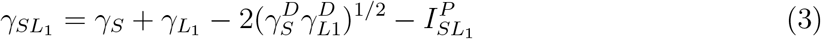

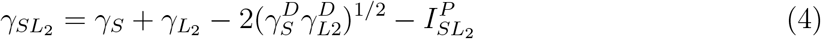

A single letter subscript refers to the surface tension of the prescribed phase. A double letter subscript refers to the interfacial tension between two phases. The superscript, *D*, refers to the dispersive contribution driven by London dispersive force, and the superscript, *P*, refers to the polar contribution driven by hydrogen bond and other polar interactions.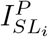refers to the polar interaction between solid substrate (cell) and liquid phase. Here we recognize that 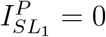 in Eq. 3 as non-polar liquid does not exhibit polar interactions.

Young’s equation at the solid-liquid-liquid interface is,

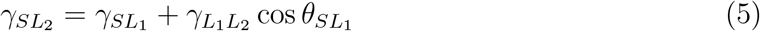

Substituting the Fowkes expressions for interfacial tensions (Eq.3-4) into Young’s equation (Eq.5) yields the following expression,

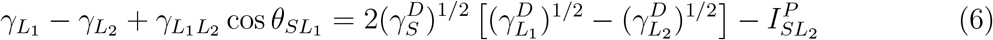

Eq.6 states a linear relation between the independent variable, 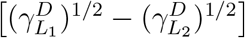, and the dependent variable, 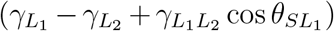. The surface tension of non-polar liquids were collected from previous literature and reproduced in Table **??**. The surface tension of the aqueous phase was measured by pendent drop method. A linear regression was done with a series of *n*-alkane liquid phases, in which the slope signifies 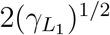, and the intercept signifies 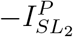.

## 6 Acknowledgement

The authors would like to acknowledge the Dunn Lab at Stanford University for providing cell culture facilities and fruitful discussions. C.L. was supported by a Stanford Bio-X fellowship. The delamination experiments were performed at the Stanford Nano Shared Facilities (SNSF), supported by the National Science Foundation under award ECCS-1542152.

