## Supplementary Materials for "Air-Liquid Interface Induced Epithelial Delamination"

#### Author Information

**Chunzi Liu**
Department of Chemical Engineering, Stanford University Stanford, CA 94305

**Gerald G. Fuller**
Department of Chemical Engineering, Stanford University Stanford, CA 94305


### 1 Supplementary Methods

#### Measure fluid phase surface tension

To determine the aqueous phase surface tension, $\gamma_{LV}$, an air bubble of 2 $\mu$L was generated with a Ramé-Hart 290 contact angle goniometer in an aqueous (phosphate buffer solution) environment in the presence of ocular epithelium. Ten images were taken in a quick succession. The shape of the air bubble was analyzed using DROPimage Advanced, a software developed by Ramé-Hart. The surface tension was then calculated using the Laplace-Young equation. To determine the interfacial tension between $n$-alkane phase and the aqueous phase, a similar method was employed in which an alkane droplet was generated in an aqueous environment as shown in Fig S1.

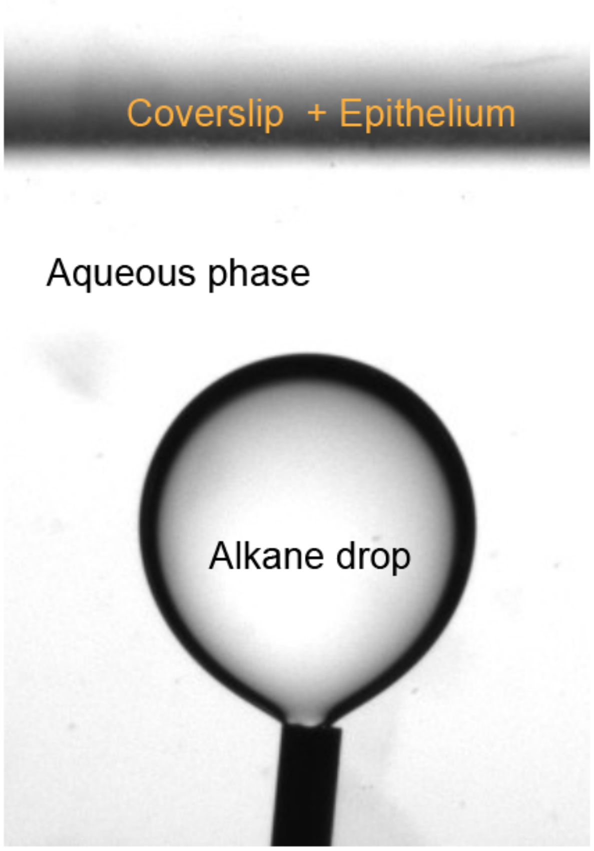


Figure 1: An example image to measure the alkane surface tension.

### 2 Supplementary Table

The following table lists the surface tension and interfacial tension against water for alkanes used in measurements.

Table 1: Surface tension and interfacial tension of linear alkanes

| Alkane | $\gamma_H$ | $\gamma_{WH}$ |
|---|---|---|
| n-hexane | 18.43 | 50.80 |
| n-heptane | 20.14 | 51.24 |
| n-octane | 21.60 | 51.64 |
| n-decane | 23.83 | 52.33 |
| n-octane | 25.35 | 52.87 |
| n-octane | 27.47 | 53.50 |

Alkane surface tension values are retrieved from http://www.surface-tension.de/. Alkane interfacial tension values are retrieved from internal database from DataPhysics Instruments USA Corp which was requested via.

### 3 List of Movies

All movies will be shared upon request from the authors.

Movie S1. Ocular epithelium exhibiting static behavior

Movie S2. Ocular epithelium exhibiting delamination and recovery behavior

Movie S3. Ocular epithelium exhibiting rupture behavior

Movie S4. Ocular epithelium under EDTA and latrunculin-A treatment.
